# Impact of Cervical Lymphatic Obstruction on Brain Pathophysiology in Cervical Lymphedema Animal Models

**DOI:** 10.1101/2024.02.21.581490

**Authors:** Hwayeong Cheon, Dong Cheol Woo, Seungwoo Cha, Yeon Ji Chae, Inhee Maeng, Seung Jae Oh, Jae Yong Jeon

## Abstract

**Background:** Injury to the cervical lymph nodes can lead to cervical lymphedema and subsequent fluid accumulation in the head and neck region, potentially causing pathophysiological alteration in the brain. This condition is thought to be linked with various neurological diseases, although the direct connection between cervical lymphatic obstruction and its effect on the brain has been difficult to establish.

**Methods:** We produced the disease animal models through lymph node dissection and radiation in fifteen male Sprague–Dawley rats aged 8 weeks and weighing 280–320 g. The models were specifically designed to induce lymphatic obstruction in the cervical region only, with no direct interventions applied to the brain. We evaluated swelling and lymphatic drainage in the head and neck for follow-up. The size of the lateral ventricles was verified through MRI, and changes in water content in brain tissue were directly measured. At 2 and 8 weeks, we observed immune cell infiltration, ventricular enlargement, and pathohistological changes in the harvested brain tissues.

**Results:** The experimental animals exhibited lymphatic obstruction in the cervical region, with swelling, abnormal lymphatic drainage, and immune cell infiltration into the brain’s white matter, reminiscent of extremities lymphedema. MRI revealed lateral ventricular enlargement in these animals, indicative of increased cerebrospinal fluid levels compared to the control group. This increase in cerebrospinal fluid was associated with an increase in brain tissue water content, leading to pathophysiological changes akin to those seen in hydrocephalus and cerebral edema.

**Conclusion:** The outcomes in this study underscore a significant link between lymphatic circulatory dysfunction and the onset of neurophysiological diseases. Cervical lymphedema showed pathophysiological changes similar to those seen in extremities lymphedema. However, these changes in the brain could be more critical than in the extremities. Our finding highlights the importance of understanding lymphatic system health in preventing and managing neurological conditions.

## INTRODUCTION

Cervical lymph nodes (CLNs) are the dominant lymph nodes into which lymph is collected in the head and neck lymphosome^1^. CLNs are the major drainage routes for cerebrospinal fluid (CSF) flowing through the nasal cavity into the lymphatic system, highlighting the importance of CLNs in fluid circulation and immune function^2–6^ (Figure S1A). They play an important role in the head and neck by filtering and transporting the lymph from the CSF. Moreover, a recent study reported that lymphatic vessels at the back of the nose serve as a primary outflow pathway of CSF to CLNs, suggesting a close relationship between the function of CLNs and drainage of CSF^7^ (Figure S1B). Therefore, CLN dissection for head and neck cancer treatment can lead to lymphatic obstruction in the cervical region, thereby increasing CSF levels in the central nervous system (CNS). The recent discovery of lymphatic circulation in the CNS has sparked interest in its role in fluid balance, waste clearance, immune response regulation, and neurological disease pathogenesis^8–11^. Several studies on CNS lymphatics have focused on the onset of Alzheimer’s disease, which is associated with beta-amyloid excretion in relation to aging^12–15^. However, irrespective of beta-amyloid excretion, lymphatic obstruction in the head and neck region may involve another mechanism that can lead to other neurological diseases. Lymphatic circulation disorders and the resulting disruption of water balance are major causes of lymphedema, and the occurrence of lymphedema leads to immune cell infiltration and inflammation at the affected site^16,17^. Since the lymphatic obstruction in head and neck can lead to an increase in water content and immune cell infiltration in the brain, cervical (head and neck) lymphedema could be a cause of neurological diseases distinct from Alzheimer’s disease. In clinical settings, lymphedema occurs when the dominant lymph nodes are damaged or removed because of lymphatic injury, such as lymph node dissection. Therefore, we hypothesized that CSF accumulation caused solely by CLN dissection may lead to pathophysiological and histopathological alterations in the brain, potentially triggering neurological disorders. (Figure S1C).

To validate our hypothesis, we produced animal models with cervical lymphedema or lymphatic obstruction in the head and neck region via superficial and deep CLN (SCLN and DCLN) dissection and radiation to investigate pathophysiological alterations in the brain. Therefore, complete removal of the SCLN and DCLN is likely to induce cervical lymphedema. We then used preclinical magnetic resonance imaging (MRI) and terahertz (THz) spectroscopic imaging systems to measure the increase in CSF and water content in the brain in the animal models of lymphedema. We investigated how the observed increase in CSF and water content in the brain tissue induces pathological changes and identified similarities between these pathological changes and existing neurological disease alterations.

## METHODS

All experiments were conducted in accordance with the relevant guidelines and regulations of the Institutional Animal Care and Use Committee (IACUC) of our institution (2022-12-110 and 2023-30-226). The IACUC abides by the Institute of Laboratory Animal Resources and Animal Research: Reporting of *in vivo* Experiments guidelines of the National Center for the Replacement, Refinement, and Reduction of Animals in Research.

### Production of animal models of cervical lymphedema

All animals were given free access to water and food and were kept under stable humidity and temperature. This study included 15 male Sprague–Dawley rats weighing 280–320 g (8 weeks old), and 12 animals were the experimental group (CLN dissection group), while 3 animals were in the control group. The rats were fed adaptively for 1 week before surgery, and all procedures involving animal experiments were conducted in specific designated areas. The development of animal models of lymphedema for preclinical research is still an actively researched area, but the fundamental surgical and radiation protocols followed the methods used for producing extremity lymphedema animal models^59–62^ based on the research by Daneshgaran et al.^63^. Before surgery, rats were anesthetized with a mixture of tiletamine/zolazepam (Zoletil 50) (50 mg/kg; Virbac, France) and xylazine (Rompun, Bayer Korea, Seoul, Republic of Korea) at a volume ratio of 5:1 after induction with 4% isoflurane gas. During anesthesia, electric clippers were used to remove fur from the cheek and neck.

All animals were operated on by the same investigator. After disinfecting the surgical area with 75% ethyl alcohol, microsurgical procedures for lymph node dissection were performed. A circumferential skin incision was made on the ventral side of the neck. Subsequently, 0.05 mL Evans blue (Sigma Aldrich Co., MO, USA) solution (30 mg/mL in 0.9% saline) was subcutaneously injected into the lower lip and both ears, enabling CLN visualization. SCLNs are circularly arranged around the submandibular glands of the masseter muscle and are connected by collecting lymphatic vessels that extend from the cervical region^64^. SCLNs (lymph nodes 1–3 in Figure S2A) are sequentially connected to DCLNs along the lymphatic vessels that collect lymph (lymph node 4 in Figure S2A). After visualization using Evans blue solution, the lymph nodes were carefully removed from the deep surface with subcutaneous fat to minimize iatrogenic damage. Given that the nerve plexus, major veins, and arteries are distributed around the CLNs, we exercised caution during incision and cauterization to avoid damaging the adjacent tissues (Figure S2B).

After the procedure, the skin edges were sutured using the folding suture method, bringing the skin surfaces into contact with each other to prevent intradermal lymphatic vessel reconnection. The skin incision was circumferentially cauterized before suturing. Ketoprofen (1 mg/kg; SCD Ketoprofen Inj., SamChunDang Pharm, Seoul, Republic of Korea) was administered intramuscularly immediately after the surgery. On postoperative day 2, the surgical site, including the neck area, was irradiated with a single dose of 20 Gy using an X-Rad 320 device (Precision X-Ray Inc., CT, Madison, WI, USA). The remaining parts of the body including the head were shielded with customized 8-mm thick lead plates (99% radiation shielding) to protect them from radiation exposure. Animals were anesthetized with 4% isoflurane gas before irradiation and placed in the prone position. A cumulative radiation dose of 20 Gy was delivered in 10 fractions at a rate of 1 Gy/min. Only animals with successful model formation were used for the experiment 1 week after radiation exposure, while the rest were sacrificed. Model formation was determined by evaluating edema and lymphatic drainage at 1 and 8 weeks after surgery and radiation.

### Cervical edema evaluation and NIRF-ICG lymphangiography

In previous studies, the development of lymphedema in animal models was verified via limb circumference or thickness measurement^65,66^. In this study, changes in the cervical cross-sectional area were used to evaluate edema. We estimated the cervical cross-sectional area based on the anatomy of the rats, assuming an elliptical shape with the vertebra in the middle. Lateral and ventral images of the neck were consistently measured and analyzed from the same posture, and the cross-sectional area was calculated according based on the neck diameter using ImageJ software (ImageJ 1.48 v, http://rsbweb.nih.gov/ij/; NIH, Bethesda, MD, USA). After anesthesia, the animals were flexed on a plane in a completely relaxed state, and the diameter of the neck was measured using a customized near-infrared fluorescence (NIRF) imager (Figure S3)^18,61,67,68^.

Near infrared fluorescence indocyanine green (NIRF-ICG) lymphangiography is the primary technique for evaluating lymphatic drainage, allowing real-time imaging of lymphatic function and structure. We employed NIRF-ICG imaging to determine the condition of cervical lymphatic drainage^69^. As hair scattered infrared light, the animals were sedated using 4% isoflurane gas, and the hair on the region of interest was removed using electric clippers and depilatory cream to facilitate an accurate observation of fluorescence images. Subsequently, 3 μL of indocyanine green (ICG) powder (Diagnogreen Injection 25 mg; Daiichi Sankyo co., LTD, Tokyo, Japan) mixed 25 mg/ml bovine serum albumin (BSA) solution (20 μg/ml; Sigma, St. Louis, MO) was intradermally injected into the lower rip using 34-gauge needles. In clinical practice, abnormal lymph drainage patterns are detected using NIRF-ICG lymphangiography in patients with lymphedema. An ICG agent injected into the lymphatic system can be detected because it emits fluorescence when excited by an external near-infrared light source. Based on the clinical standards for lymphedema severity, abnormal lymphatic drainage patterns are categorized as splash, stardust/diffuse, or blackout. In normal lymphatic circulation, most of the lymph collects and flows through collective lymphatic vessels in a linear pattern. The splash pattern represents dermal backflow in the superficial pathway. Stardust/diffuse patterns caused by lymphatic accumulation and leakage indicate severe lymphedema. Finally, the blackout pattern indicates the absence of lymphatic drainage due to obstruction.

Figure S4 shows that severe lymphedema also resulted in an increased flow of lymph into the surrounding tissues, thereby causing the linear ICG fluorescence image to become planar. Given that such changes increase the bright area (area where contrast agent spread) in the images, the severity of lymphedema can be measured by calculating the area. In this experiment, we measured the bright area using the automatic threshold function of ImageJ software. The images were acquired at a constant height of 25 cm from a fixed position. We quantified the amount of lymph flowing into the surrounding tissues using NIRF-ICG lymphangiography (Figure S5). In addition, we measured lymphatic contraction signals in the same collective lymphatic vessels in both groups. In a previous study, we examined a method for obtaining lymphatic contraction signals through signal processing and identified variations in lymphatic contraction signals caused by lymphatic obstruction^18^. Lymphatic obstruction in the animals used in this study was evaluated using these variations in lymphatic contraction. Therefore, we obtained 15-min real-time videos to identify these variations in lymphatic contraction and extracted lymphatic contraction signals in both groups using ImageJ software. FFT signal processing was performed using Origin Pro 9 software (Origin 9.0; OriginLab, Northampton, MA, USA) for numerical calculations and visualization based on previous research^18^.

### Volumetric analysis of the brain using MRI

In the animal models, an increase in CSF levels in the brain with lymphedema was verified 2 and 8 weeks after surgery and radiation using high-precision MRI. In the CNS, the CSF flows through a system of fluid-filled cavities known as the ventricular system. CSF drainage disturbance increases CSF pressure within the ventricular system, leading to an abnormal increase in the ventricular system volume ^70,71^. To measure the volume of the ventricular system, MRI was performed using a 7.0 T Bruker preclinical MRI scanner (PharmaScan 70/16; Brucker BioSpin GmbH, Germany) with a 72-mm transmit volume coil and a rat brain surface receiver coil. All the animals were anesthetized with 1.5–2.5% isoflurane in a mixture of 70% nitrous oxide and 30% oxygen administered through a nose cone. A warm-water circulating flatbed was used to maintain their body temperature at 37.5 ± 0.5°C, and they were consistently monitored for stable breathing. T2-weighted images (T2-WIs) were acquired using a fast spin-echo sequence (TR, 5250 ms; TE, 66 ms; averages, 1; echo spacing, 11 ms; field of view, 25 × 25 mm; slice thickness, 1.0 mm; slice number, 27). MRI images were analyzed using ImageJ software. Herein, the total brain and lateral ventricle (LV) volumes were measured using T2-WIs and the two-dimensional volumetric technique, which involves adding the volume measured from each slice.

### THz spectroscopic imaging system

The THz spectroscopic imaging system used to measure the brain tissue water content consisted of an ultrashort pulsed laser with a center wavelength of 1560 nm and two fiber-coupled antenna modules (FC/APC) for the emitter/detector. For the laser radiation, the repetition rate was 100 MHz, and the pulse width was 60 fs. The laser generated approximately 60 mW of power at the output of the polarization-maintaining fiber. The laser beam was divided by a beam-splitter fiber and guided to the emitter and detector. The emitter antenna (Tx; TERA 15-TX-FC; Menlo Systems GmbH, Planegg, Germany) had a line pattern on a Fe: InGaAs/InAlAs substrate and was equipped with a PM PANDA fiber patch cord (I = 100 cm, FC/APC connector). THz waves were generated from the emitter with an 80-V DC bias. The THz waves were incident vertically on two polymethylpentene (TPX) lenses (L1 and L2) on the sample holder by a metal mirror (M1), reflected off the sample, and guided into the detector by a Si beam splitter (THz BS) and TPX lens (L3) (Figure S6). The detector antenna (Rx; TERA 15-RX-FC; Menlo Systems GmbH, Planegg, Germany) had a dipole pattern on an LT InGaAs/InAlAs substrate with the same fiber patch cord as the emitter. THz pulses were acquired by the pump-probe sampling method using a fast scan delay with a resonance frequency of 20 Hz and a scan range of 30 ps. The THz pulses were recorded automatically by a computer using a low-noise current preamplifier and data acquisition board. THz spectroscopic images of the sample with a spatial resolution of 250 × 250 μm per pixel were obtained using two-dimensional raster scanning. We extracted THz spectroscopic images at this frequency using the FFT of the time-domain signals. The samples were placed on a quartz window with a diameter of 30 mm and thickness of 3 mm.

### THz spectroscopy for brain sections

Eight weeks after surgery and radiation, brain samples, including the cerebrum and cerebellum, were harvested, and the weight of the brain without the meninges was measured. The brain tissues, including the cortex, white matter tract, hippocampus, and thalamus, were subsequently sectioned. The location of the sections was determined by referencing section levels 2 and 3 from the study by Rao et al. or section levels 3 and 4 from the study by Garman et al.^23,24^. The section levels were labeled based on the findings of Garman et al. After slicing in the coronal direction of the brain at section levels 3 and 4, based on the study by Garman et al.^23,24^, THz spectroscopic images were measured as quickly as possible. The sample holder containing the brain sections was sealed during scanning to prevent the tissues from drying. The samples were placed on a 2-inch quartz window for measurements using the THz spectroscopic imaging system in reflection mode. The window containing the brain tissue was sealed in a small box to prevent the sample from drying out. The sample holder was connected to two high-resolution mechanical stages, allowing for two-dimensional movement (horizontal x-y plane). This enabled images to be obtained using raster scanning with 250-μm spatial resolution. The THz spectroscopic images were analyzed separately for peak-to-peak and individual frequency-based images.

### Histological analysis

At 8 weeks postoperatively, the animals were euthanized using carbon dioxide asphyxiation, and whole brain tissues were harvested and preserved in 10% buffered formalin for 24 h at 4°C. This was followed by a minimum of 48 h of decalcification using Calci-Clear Rapid (National Diagnostics, Atlanta, GA, USA) histological decalcifying reagent. Brain sections were also taken at the same location as the THz spectroscopic imaging sections. The samples were embedded in paraffin blocks and then sectioned into 3-µm-thick slices before staining with hematoxylin and eosin (H&E). The tissue sections were examined using a 20× objective microscope equipped with an integrated camera (Olympus B53, Tokyo, Japan). Tissue density was determined by scanning and digitally processing H&E-stained slide images. All brain sections were acquired for comparison with THz spectroscopic images. All images were processed and assessed using Olympus Cellsens Standard ver. 1.13 software (Olympus, Tokyo, Japan). The following antibodies were used for immunohistochemical staining of brain sections: rabbit monoclonal CD11c C-terminal (1:200; ab52632, Abcam, Cambridge, UK) as a DC marker and rabbit monoclonal CD4 (1:200; ab133616, Abcam, Cambridge, UK), which is a T-cell receptor marker. All slides were imaged using a light microscope (Model BX40, Olympus, Tokyo, Japan). During imaging, the cell nucleus area was measured and quantified by selecting three regions from brain sections prepared from three rats in each group using the color threshold function of ImageJ software.

### Statistical analysis

All the statistical analyses were performed using GraphPad Prism 9 (GraphPad Software Inc., Boston, MA, USA) and Microsoft Excel 2019 version 2111 (Microsoft Corporation, Redmond, WA, USA). Student’s t-tests were used to compare differences between groups. A *p*-value < 0.05 was considered to indicate a statistically significant disparity.

## RERULTS

### Volume evaluation in the cervical region

Edema or swelling is the most common diagnostic criterion for lymphedema. In the animal models used in this study, edema was observed in the cervical region and cheeks 1 week after CLN dissection and radiation and swelling was consistently maintained during the follow-up period (Figure S7). To quantify the degree of edema, we compared the CLN dissection group with the control group. The CLN dissection group showed an approximately 30% increase in neck cross-sectional area compared with that in the control group (*p* < 0.05) (Figure 1). This indicated that significant cervical lymphedema occurred in the CLN-dissected animal model.

**Figure 1.**
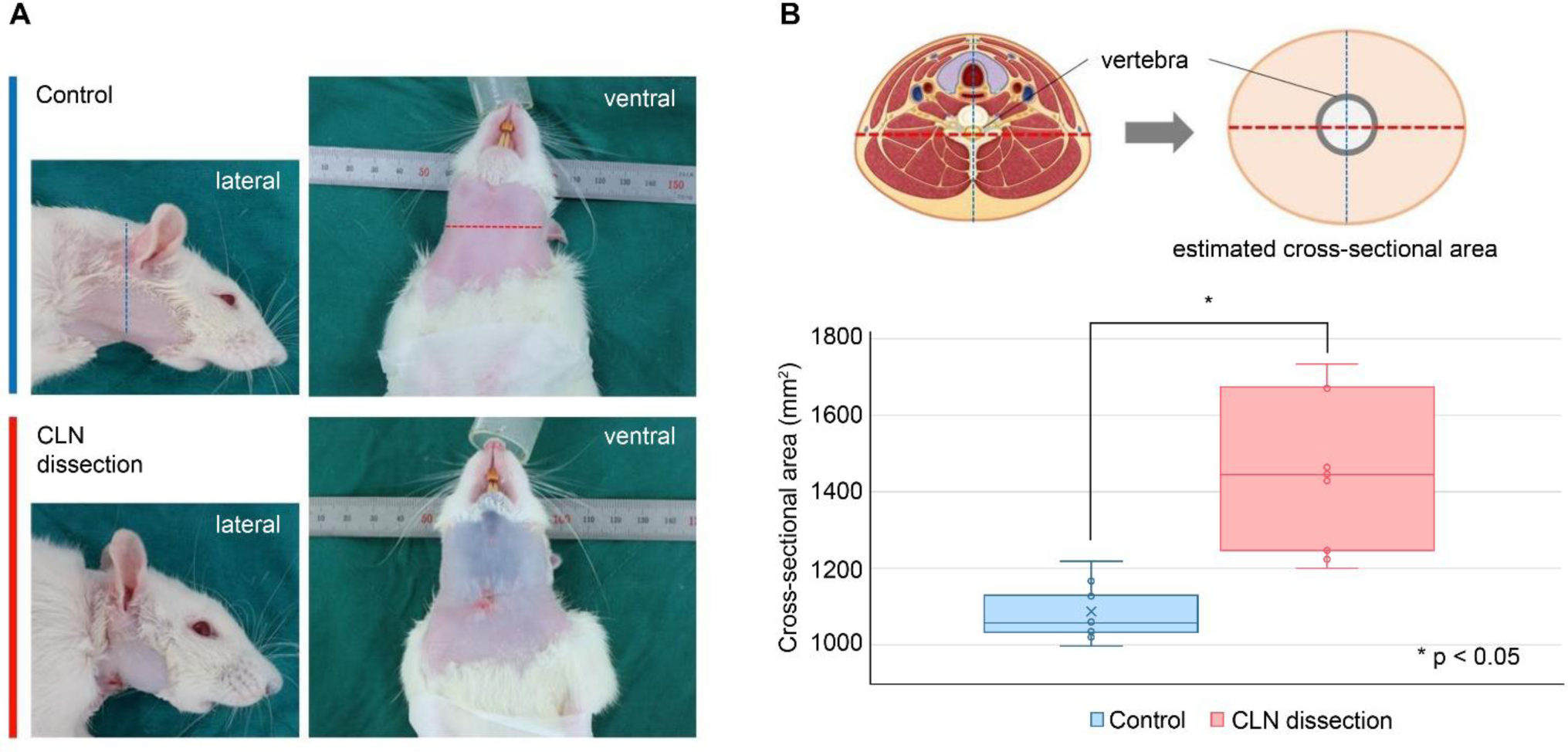
Measurement of neck circumference changes following cervical lymph node (CLN) dissection. (a) Lateral and ventral views in the cervical region for the control and CLN dissection groups. (b) Neck cross-sectional area for the control and CLN dissection groups. The shape of the neck cross-section is assumed to be an ellipse centered on the vertebra, based on cervical anatomy. The cross-sectional area differs significantly between both groups (Student’s t-test, *p < 0.05).

### Evaluation of CLN obstruction and increased CSF levels

Subsequently, we verified whether the edema observed in the animal models resulted from lymphatic obstruction due to CLN dissection. In the control group, lymphatic flow occurred in the most rostrally located SCLNs along the collective lymphatic vessels in the cervical region before draining into subsequent CLNs. However, in the CLN dissection group, abnormal lymphatic drainage patterns (splash and stardust) were observed throughout the ventral cervical region, including the neck and face. Lymph gradually flowed along the net-shaped (splash-patterned) superficial lymphatic vessels. However, dermal backflow due to obstructed lymph drainage was observed (Figure 2A). The CLN dissection group had a significantly higher degree of lymph drainage than the control group (Figure 2B) (*p* < 0.05).

**Figure 2.**
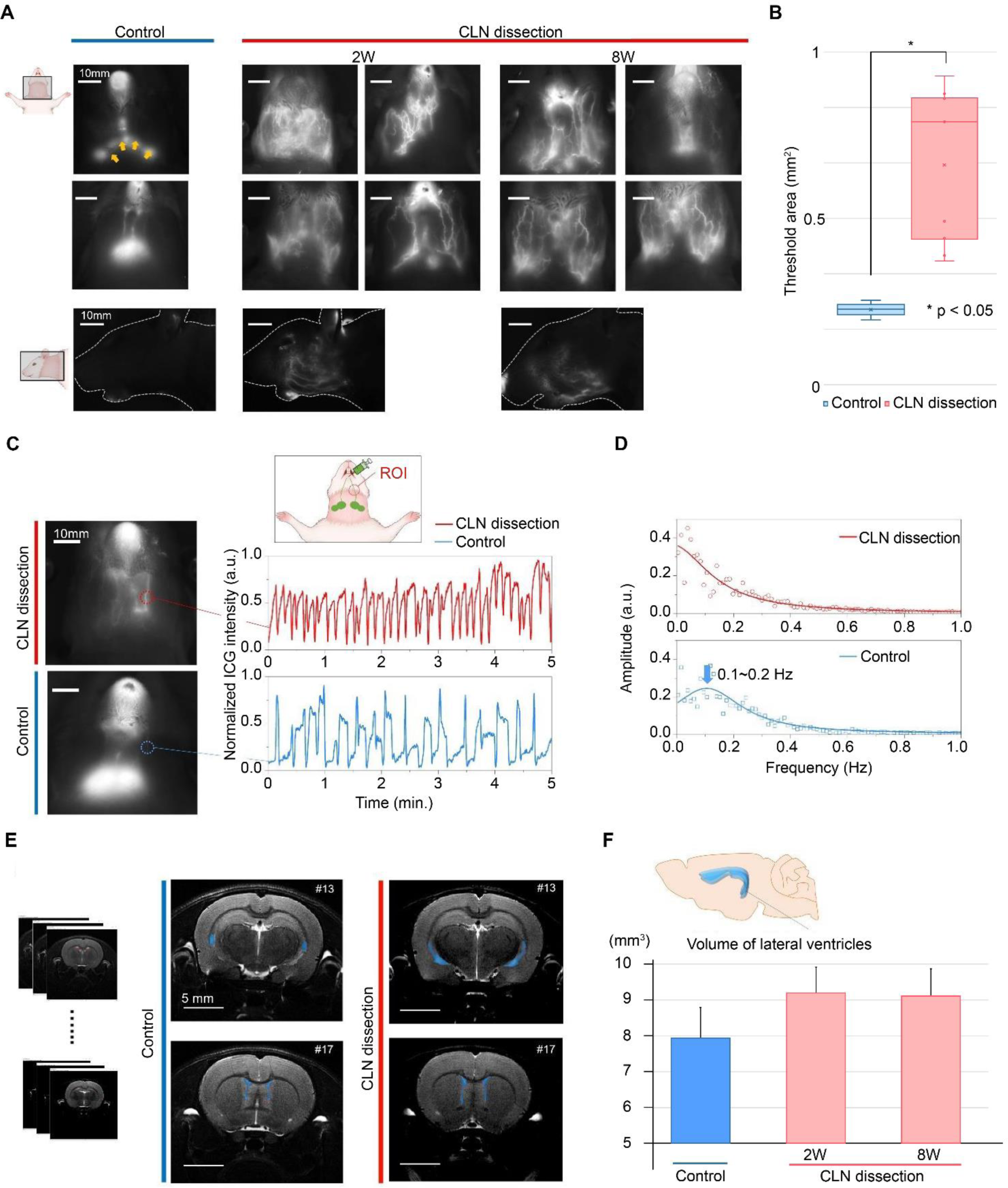
Measurement of lymphatic obstruction using near-infrared fluorescence indocyanine green (NIRF-ICG) lymphangiography and cerebrospinal fluid (CSF) level increase using magnetic resonance imaging (MRI). (A) Lymphatic drainage in the cervical region of control and cervical lymph node (CLN) dissection group animals (at 2 and 8 weeks) using NIRF-ICG lymphangiography. In the control group, the lymph flows along the collective lymphatic vessels toward the CLNs (yellow arrows). In the CLN dissection group, the CLNs are not found, and abnormal lymphatic drainage patterns are observed in the neck and face at 2 and 8 weeks after surgery owing to lymphatic obstruction. (B) Lymph flow quantification. The threshold area is significantly different between the groups (Student’s t-test, \**p* < 0.05). (C) The time-domain waveforms of lymphatic contractions is measured in the regions of interest (ROI, red dotted circle) of each group. Compared with the control group, the waveform and frequency of lymphatic contractions are alternated in the CLN dissection group. (D) Signal conversion from the time-domain to the frequency-domain spectrum via Fast Fourier transform signal processing. In the control group, signal peaks are observed between 0.1 and 0.2 Hz, indicating frequent movements with a consistent period. (E) The images of # 13 and 17 slides among 27 magnetic resonance imaging (MRI) sections are used to measure the volume of the lateral ventricle in the control and CLN dissection groups. The area within the red line represents the lateral ventricle. (F) The volume of the lateral ventricle was determined by MRI in the control and CLN dissection groups at 2 and 8 weeks after CLN dissection. The volume of the ventricle in the CLN dissection group is increased by approximately 15% compared with that of the control group (at 2 and 8 weeks), indicating an increase in CSF levels.

Furthermore, abnormal waveforms of lymphatic contraction were observed in the CLN dissection group. A previous study showed a change in lymphatic contraction induced by lymphatic obstruction in upper limb lymphedema models^18^. Figure 2C demonstrates a disruption of the cervical lymphatic circulation. In the CLN dissection group, we observed a waveform change and an increase in the frequency of lymphatic contractions compared with those in the control group. The control group presented a more robust lymphatic contraction frequency than the CLN dissection group, which was verified by transforming the time-domain data in Figure 2C into a frequency-domain spectrum through Fast Fourier transform (FFT) signal processing. The control group showed regular lymphatic contractions with a peak frequency between 0.1 and 0.2 Hz (Figure 2D). In contrast, the CLN dissection group showed irregular contractions owing to the disturbances caused by these robust movements.

Volume measurements and near-infrared fluorescence indocyanine green (NIRF-ICG) lymphangiography demonstrated cervical lymphatic obstruction in CLN-dissected animals. Owing to the strong correlation between cervical lymphatic flow and CSF drainage, we investigated whether cervical lymphatic obstruction increases CSF levels. In the animal models, MRI was used to compare the LV volume change in the CLN dissection and control groups (Figure 2E). The LV volume in the CLN dissection group increased by approximately 15% compared with that in the control group at 2 weeks and 8 weeks postoperatively (Figure 2F). Furthermore, in the CLN dissection group, the LV to whole brain volume ratio increased from 3.8% to 4.7% and 4.4%. MRI results indicated an increase in CSF levels in the CLN dissection group, suggesting a corresponding increase in brain tissue hydration.

### Evaluation of brain water content using THz spectroscopic imaging

CLN dissection-induced lymphatic obstruction was identified based on edema and abnormal lymphatic drainage in the cervical region. MRI confirmed increased CSF levels in the brain due to lymphatic obstruction. Therefore, an increase in brain water content may have resulted from CSF accumulation. We observed changes in the hydration of freshly excised brain tissues using THz reflectance in THz spectroscopic imaging. A previous study measured the water content of animal brain tissue using THz spectroscopy and verified its accuracy by measuring the tissue water mass, confirming that THz spectroscopy can precisely measure the water content in biological tissues^19^. Considering that important pathophysiological changes often lead to increased tissue water content, which may serve as a diagnostic criterion for specific diseases^20–22^, THz spectroscopic imaging could be a valuable tool for detecting brain tissue water content.

Before THz spectroscopy, the entire brain was weighed after harvesting from the animal (Figure 3A). The brain weight in the CLN dissection group was approximately 11% higher than that in the control group; however, the difference was not statistically significant. Figure 3B shows reflectance images of the brain tissue sections of level 3 and 4 obtained using THz peak-to-peak values. The sectional location was based on references to previous researches^23,24^. The CLN dissection group showed a higher proportion of red and white areas, indicative of a higher water content within the tissue, than the control group. The individual frequency imaging at 0.5, 1.0, and 1.5 THz showed that the 0.5 THz reflectance of brain tissue in the CLN dissection group was significantly higher than that in the control group (Figure S8). This result indicates that our THz spectroscopic images detected an increase in brain tissue water content. This is attributed to the sensitivity of the lower-frequency bands in the THz range, which can detect more changes in water content caused by the higher refractive index of water molecules^25–27^. Based on the results of the individual frequency images, the difference between the two groups in the THz spectroscopic images was derived from the difference in THz reflectance caused by the brain tissue water content. The spectral images were reconstructed into binary THz spectroscopic images based on higher (red) and lower (blue) water contents (Figure 3C). Although high water content was only detected around the LV in the control group, we observed an increase in water content throughout a wider area of brain tissue in the CLN dissection group. These changes were more distinct in the level 4 section than in the level 3 section. In the binary images, the red areas indicate regions with an average increase in water content of approximately 5-6% compared with the blue areas based on individual frequency imaging and the integral values of the spectrum at each pixel point (Figure 3D).

**Figure 3.**
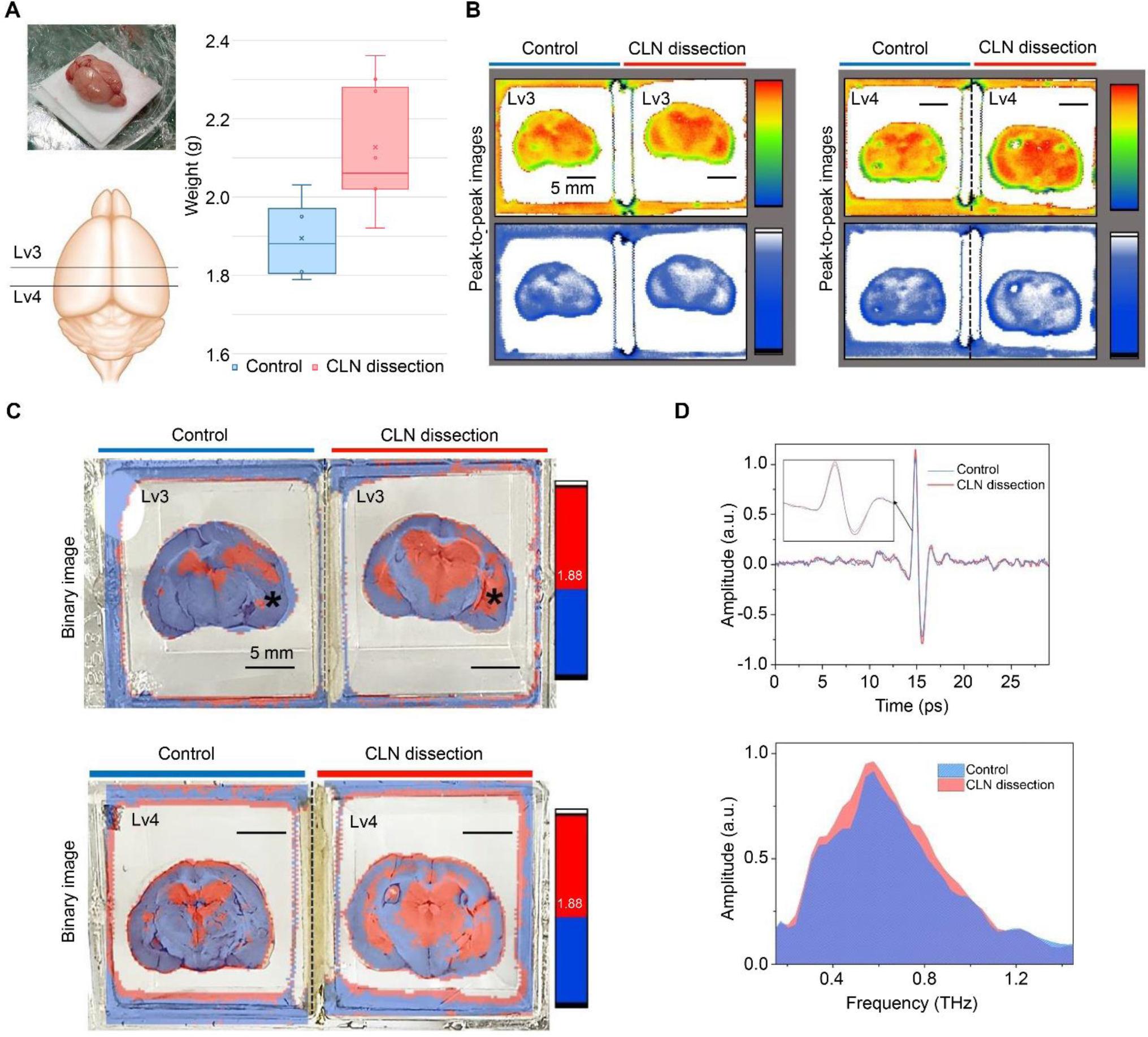
Terahertz (THz) imaging of brain changes following cervical lymph node (CLN) dissection. (A) The brain section locations of levels 3 and 4 and the weight of the whole brain are measured immediately after extraction from the animals in each group. (B) The peak-to-peak THz reflectance images of the brain sections are represented by the rainbow and blue-white gradual values. (C) The overlay visible light imaging combined with THz reconstructed binary imaging (higher level: red, lower level: blue) of sectional brain sections. The CLN dissection group at 8 weeks shows a widely higher reflectance than the control group in both sectional levels. (D) Time- and frequency-domain THz spectra of the pixels marked with black asterisks in control and CLN dissection group images of Figure 3C. All scale bars of THz images (b and c) are 5 mm.

### Histological alterations in brain tissue

The red area in Figure 4A represents the region where the water content was relatively high in THz spectroscopy, which was overlaid with the brain tissue section for pathological examination. The locations of the LV and third ventricle (TV), where the CSF accumulates, are closely associated with the distribution of tissues with increased water content. In the control group, the tissues surrounding the LV and TV were more hydrated, which was similar to the results obtained with brain section images. However, the CLN dissection group showed an increase in water content over a wide brain tissue area, including the ventricular region.

**Figure 4.**
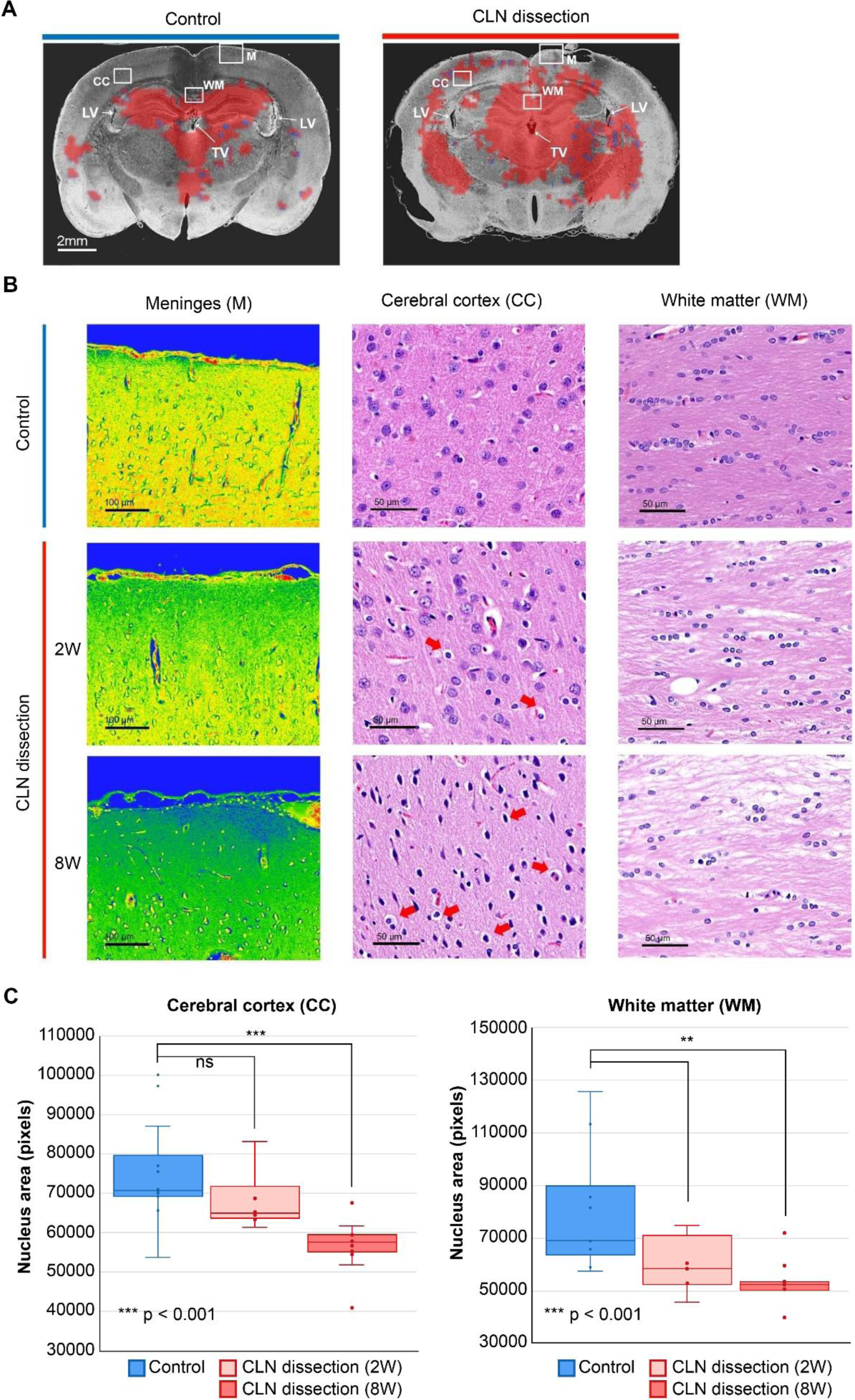
Histological brain tissue alterations in the cervical lymph node (CLN) dissection group. (A) The merged images between the region of high-water content by measuring THz imaging and fixed tissue in the paraffin block. The structural alterations in brain tissue are primarily investigated around regions with differences in water content: meninges (M), cerebral cortex (CC), and white matter (WM) of corpus callosum. LV: lateral ventricle, TV: third ventricle. (B) Tissue structure comparisons in each section. In the CLN dissection group, a decrease in peripheral brain tissue density is observed along with meningeal edema. In the cerebral cortex of CLN-dissected animals, many apoptotic cells are identified, including considerable tissue density reduction (red arrows). A decrease in the tissue capsule is observed in the WM of CLN-dissected animals. These changes are more noticeable at 8 weeks (8W) than at 2 weeks (2W). (C) Quantification of the cell nucleus areas in the CC and WM regions of animals in each group. At 8W, the CLN dissection group shows a significant difference in cell nucleus area compared with the control group (Student’s t-test, \*\**p* < 0.01 and \*\*\**p* < 0.001).

Based on these images, we investigated brain tissue changes, focusing on areas with significant changes in water content on THz spectroscopy: the meninges, cerebral cortex (CC), and white matter (WM) (Figure 4B). In the meninges, where many meningeal lymphatics are present, the CLN dissection group exhibited more partial meningeal edema than the control group. This was more visible at 8 weeks after surgery and radiation than at 2 weeks. The overall tissue density decreased, and many pyknotic nuclei and cellular edema were observed in the CC. The corpus callosum of the WM also showed changes in tissue and density. These histological changes were more prominent at 8 weeks than at 2 weeks in both the CC and WM (Figure 4C). Fig. 4c shows the quantitative results of the cell nucleus area in the CC and WM in the control and CLN dissection groups (2 and 8 weeks). A decrease in the number of cells was observed in both the CC and WM regions in the CLN dissection group compared with the control group. In addition, the CLN dissection group showed a more significant difference in the number of brain tissue cells in the CC and WM at 8 weeks than at 2 weeks (*p* < 0.001 and *p* < 0.01).

Owing to the change in periventricular water content and ventricular expansion, we also examined whether characteristic periventricular histopathological alterations were present. All structures associated with ventricles showed an overall decrease in tissue density of the periventricular white matter (PVWM) and a reduction of ventral internal capsule (VIC) structure. Specifically, the ependymal cells lining of the internal surface in the ventricular space were enlarged parallel in the ventricle area. Similar to the histological changes in Figure 5, these changes were more prominently observed at 8 weeks than at 2 weeks.

**Figure 5.**
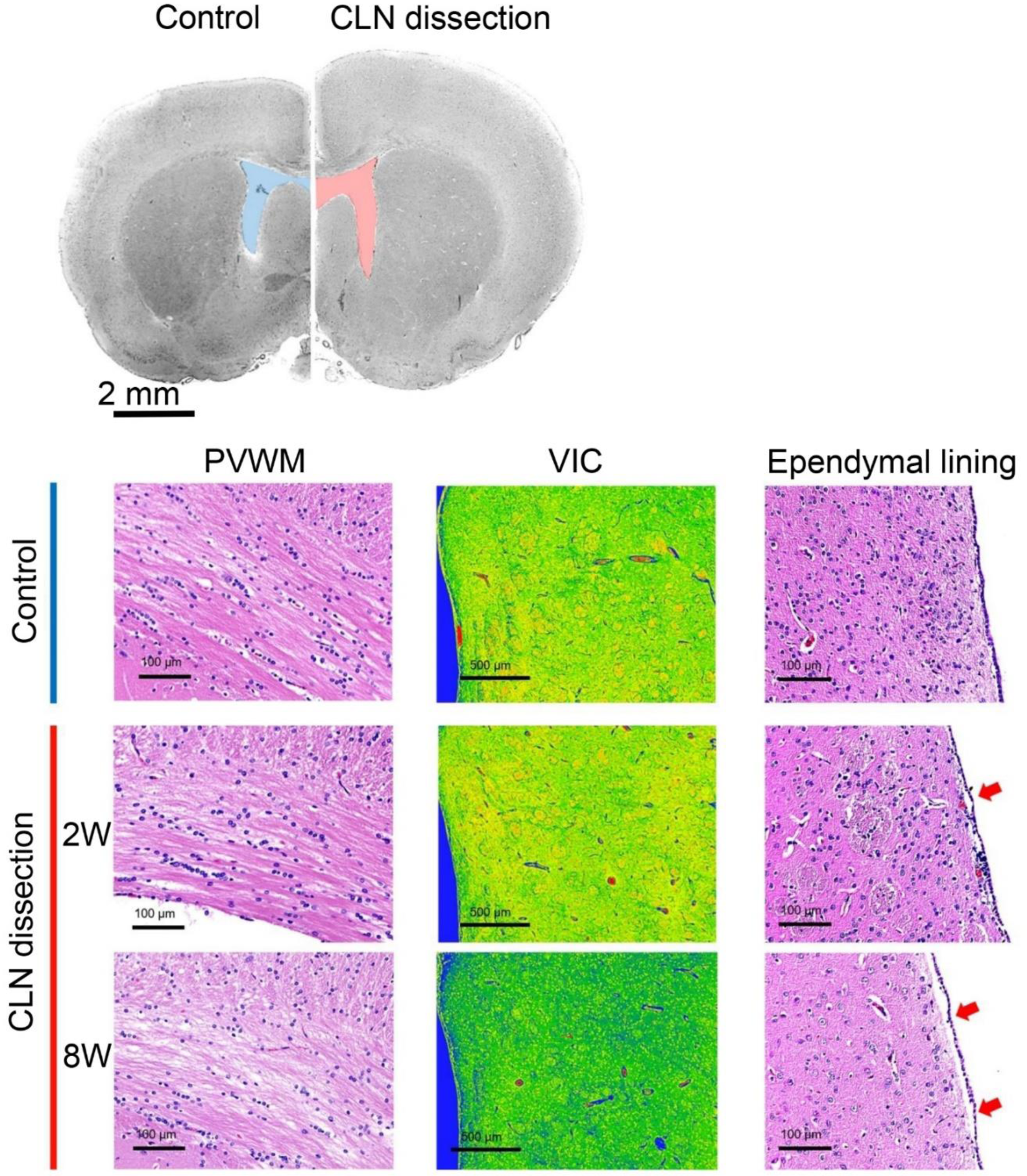
Increase in ventricular volume and associated periventricular tissue alterations in cervical lymph node (CLN)-dissected animals. In the periventricular white matter (PVWM), there is an overall decrease in density as the tissue spacing widens, and a decrease in the capsule structure of the ventral internal capsule (VIC) is identified. Additionally, an enlargement of the ependymal cell lining is observed. These changes are more pronounced at 8 weeks (8W).

### Immune cell infiltration in brain tissue

We investigated the infiltration of immune cells in the same sectional areas shown in Fig. 5 and 6 using antibodies against CD11 and CD4. As shown in Figure 6A and Figure S9, only a few immune cells were detected in the brain tissue of control group animals. It was difficult to detect dendritic cells (DCs) in most WM regions, including the PVWM, inferior colliculus, and cerebral peduncle (Figure S9). However, in the CLN dissection group, numerous immune cells were observed to infiltrate the brain tissues, and the number of infiltrated immune cells increased progressively from week 2 to week 8 in CLN-dissected animals (Figure S10). Specifically, the infiltration of DCs, a type of immune cell marked by CD11c, increased primarily around the WM region (Figure 6B).

**Figure 6.**
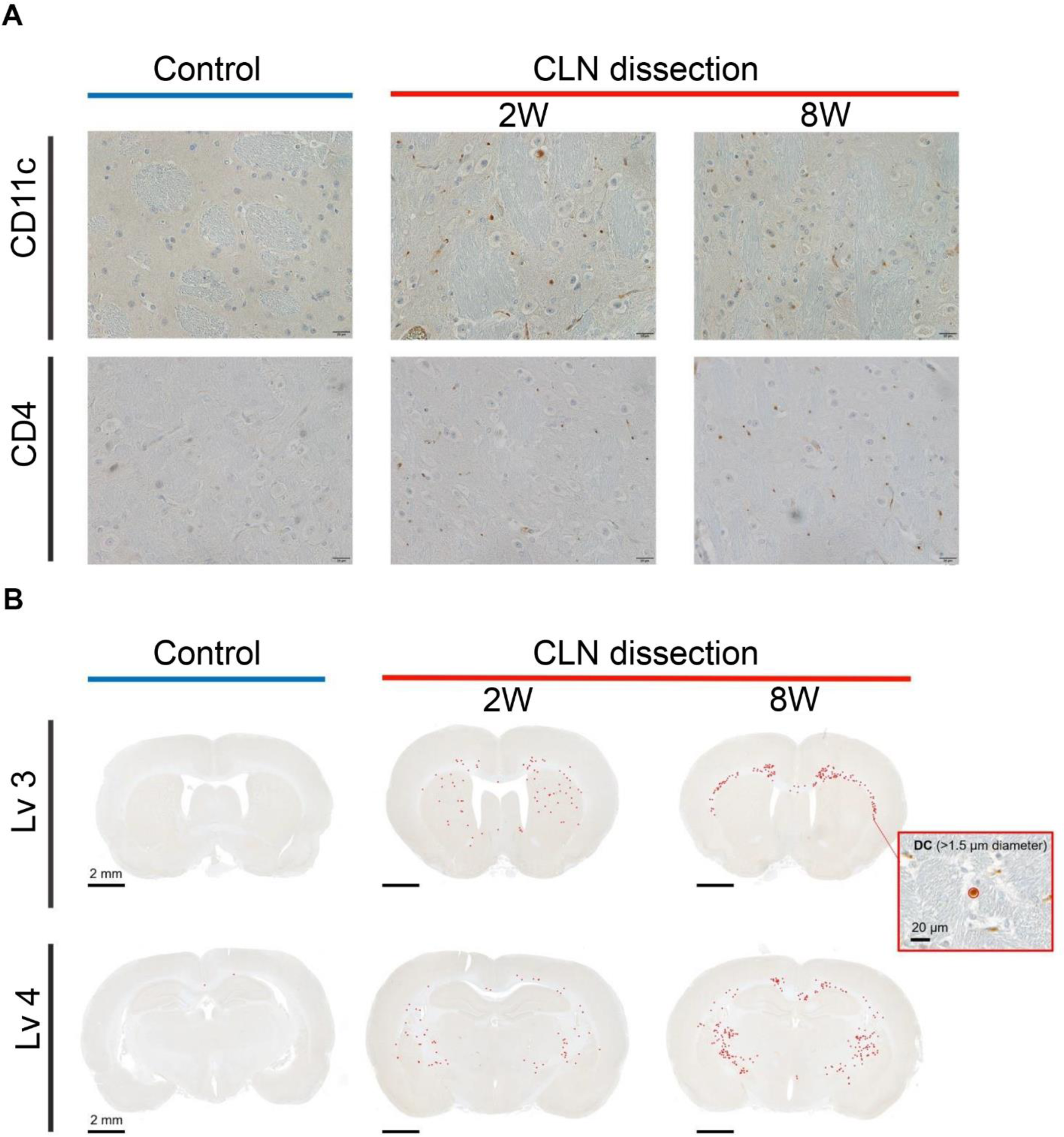
Immunohistochemical tissue examination results for the investigation of immune cell distribution in the cervical lymph node (CLN) dissection and control groups. (A) CLN dissection in animal brain sections marked with CD4 and CD11c in the ventricle-induced capsule (VIC) reveals immune cell infiltration (red arrows). The number of infiltrating immune cells increased over time from 2 weeks (2W) to 8 weeks (8W). (B) The image from each brain section represents dendritic cells (DCs) with a diameter larger than 1.5 µ***m***, marked by red circles. DC count mainly increased in the white matter.

## DISCUSSION

We investigated the increase in CSF levels within the ventricular system, increased brain tissue water content, and corresponding histological changes using animal models which dissected the bilateral CLNs and irradiated the excision sites. Although our animal model exhibited extreme lymphatic obstruction in the head and neck region compared to clinical conditions, it represents physiological conditions similar to those of patients with cervical lymphedema derived from lymph node injury. In our animal models, the increase in meningeal fluid levels was accompanied by swelling in the head and neck region (Figure S11). However, our imaging techniques enabled a more objective and quantitative assessment of these fluidic changes. MRI revealed that the LV size was further increased in CLN-dissected animals compared with control animals after 8 weeks, indicating CSF accumulation in the ventricular system. This resulted in cerebral histological changes similar to those observed in animal models of hydrocephalus. Histological examination revealed spaces between the tracts in the PVWM and a flattened lining of ependymal cells, akin to the results of a previous study^28^. Therefore, our animal model showed that hydrocephalus, ventricular enlargement, and compression of the surrounding tissues caused by abnormal CSF accumulation could be induced by lymphatic pathway disruption.

THz spectroscopic imaging at the 0.5 THz spectrum revealed an increase in water content throughout the brain tissue. The high sensitivity of the THz energy band in tissue water content measurements is a significant advantage of this imaging modality^26,29–33^. Our previous research validated that the water content analysis of the brain tissue in the 0.5-THz frequency band closely matched the water mass in the tissue^19^. These changes in brain tissue water content also led to direct histological alterations in the brain tissue. Decreased tissue density and widespread apoptotic features, such as shrunken nuclei and cellular edema, were observed in the CC of CLN-dissected animals. Given that brain tissue water content and decreased tissue density are typical manifestations of cerebral edema^34^, our animal models exhibited outcomes consistent with cerebral edema. Increased brain tissue water content is associated with cerebral edema, a critical consequence of ischemic brain injury^35,36^ and is a significant predictor of poor prognosis in neurologic disorders^37–39^. CSF is the main source of increased brain tissue water content in anoxic cerebral edema^40,41^; hence, the increase in CSF was highly associated with cerebral edema in our animal model. Importantly, these changes were attributed to lymphatic obstruction regardless of the presence of ischemic injury or stroke. Indeed, extensive cell death, shrunken nuclei, and cellular edema were observed in a wide range of brain tissue sections, consistent with previous findings in animal models of cerebral venous sinus thrombosis^42^. Similar to insufficient venous drainage, insufficient lymphatic drainage can result in cerebral edema via blood-brain barrier disruption. These changes may have substantial implications in exacerbating neuropathological disorders with potentially harmful effects on cognition.

Therefore, we demonstrated that lymphatic obstruction in the head and neck region can potentially induce neuropathological diseases, such as hydrocephalus and cerebral edema. Since the discovery of the lymphatic system of the CNS, the glymphatic and meningeal lymphatic systems have been shown to play crucial roles in the clearance of waste in the brain^43,44^. It is increasingly clear that this role is associated with the occurrence of neurological diseases, as it is related to the clearance of lethal substances that trigger neurological disorders, such as beta-amyloid^9^. Consequently, there is a demonstrated correlation between neurological disorders and the lymphatic circulation in the brain. The relationship between CLNs and CSF circulation has also been reported in various studies. Wang et al. used a DCLN model to demonstrate that the blockage of lymphatic circulation into CLNs interferes with cerebral beta-amyloid clearance, resulting in pathological features similar to Alzheimer’s disease^45^. Kinota et al. observed a blockage of CSF drainage in a DCLN ligation model using dynamic MRI^46^. Their results indicate that CLNs have a significant effect on CSF drainage. However, the relationship between cervical lymphedema-induced CSF drainage obstruction and the resulting pathophysiological and histopathological changes in the brain remains unexplored. This study revealed that blockage of the lymphatic system in the head and neck region led to the development of cervical lymphedema. Consequently, this condition results in pathohistological modifications resembling hydrocephalus and cerebral edema. We particularly consider that the alterations are associated with the inflammatory component of lymphedema.

Increased water content and subsequent alteration of tissue structures are typical histopathological characteristics, even in extremity lymphedema. Immune cell infiltration also serves as important evidence of histopathological alterations caused by lymphedema. Due to the physiological characteristics of lymphedema, this condition occurs as an inflammatory response in lymph-accumulating regions, promoting immune cell migration^47–49^. Previous studies have demonstrated immune cell infiltration in animal models of lymphedema^17,50,51^. Our study revealed immune cell infiltration in the WM of CLN-dissected animals. This feature is similar to DC accumulation in periventricular tissues due to autoimmune encephalitis, suggesting that cervical lymphedema might induce the abovementioned cerebral inflammatory responses. The infiltration of immune cells was similar to that observed in animals with extremity lymphedema^52^.

In clinical practice, lymphatic function weakening has been observed in patients who underwent chemotherapy or radiation therapy for cancer. Additionally, abnormalities in cognitive function have been reported in these patients^53–55^. Historically, edema of the neck, face, tongue, and oral cavity was considered a primary sign of cervical lymphedema in patients with head and neck cancer who had undergone CLN dissection^56–58^. However, because cervical lymphedema is also an issue of lymphatic function in the head and neck region, patients with cervical lymphedema may be at a risk of brain damage similar to that caused by the toxicity of chemotherapy and radiation. In this respect, there are limitations to this study. The bilateral CLN dissection, similar to that in the animal model used in this study, is not commonly performed in the clinical setting. Consequently, the impact on the brain may not be significant in patients with early-stage cervical lymphedema. Lymphatic circulation disorders can lead to disease-inducing changes, necessitating long-term monitoring of lymphedema, a lifelong condition. Additionally, we did not incorporate any behavioral research to assess the potential impact of these brain histopathological changes on the cognitive and motor functions of the animals. Future research should include detailed behavioral research to obtain precise results and clinical studies that track the enlargement of ventricles in patients with cervical lymphedema using MRI.

## CONCLUSION

Our study provides a compelling clue regarding the potential implications of cervical lymphatic obstruction or lymphedema on brain physiology, particularly concerning cerebral edema and hydrocephalus. These findings highlight a previously underappreciated link between the lymphatic system and neurological condition, providing valuable insights for future research. Although our study represents an important step, further investigations are necessary to fully elucidate the complex relationships between CNS lymphatic circulation and neurological diseases.

## ABBREVIATIONS

CLN: cervical lymph node
CNS: central nervous system
SCLN: superficial cervical lymph node
DCLN: deep cervical lymph node
CSF: cerebrospinal fluid
MR: magnetic resonance
THz: terahertz
NIRF: near-infrared fluorescence
ICG: indocyanine green
FFT: fast Fourier transform
LV: lateral ventricles
TV: third ventricle
H&E: hematoxylin and eosin
IHC: immunohistochemistry
DC: dendritic cell
M: meninges
CC: cerebral cortex
WM: white matter
PVWM: periventricular white matter
VIC: ventral internal capsule

## ACKNOWLEDGEMENTS

This work was supported by the National Research Foundation of Korea (NRF) grant funded by the Korean government (MSIT) (No. NRF-2019R1A2C1009055), Institute of Information & Communications Technology Planning & Evaluation (IITP) funded by the Korea government (MSIT) (Grant No. 2022-0-01044, Development of terahertz wave-based real-time intelligent brain tumor diagnosis system and technology), and a grant (2022IF0021) from the Asan Institute for Life Sciences and Corporate Relations of Asan Medical Center, Seoul, Korea.

We thank the core facilities of the Comparative Pathology Laboratory and Animal Experiment Laboratory at the ConveRgence mEDIcine research center (CREDIT), Asan Medical Center, for sharing their equipment, services, and expertise with us.

## DATA AVAILABILITY STATEMENT

The original contributions presented in the study are included in the original manuscript/supplementary material, further inquiries can be directed to the corresponding authors.

## COMPLIANCE WITH ETHICAL STANDARDS

This study was performed based on preclinical animal experiments and does not involve human-derived materials or clinical studies in the research process. The number of animals and all animal procedures in this study were approved and regulated by the Institutional Animal Care and Use Committee (IACUC) of the Asan Institute for Life Sciences, Asan Medical Center (# 2022-12-110 and 2023-30-226). The IACUC abides by the relevant guidelines including the ILAR and ARRIVE guidelines.

## AUTHORS’ CONTRIBUTIONS

H. Cheon designed the research and sample models, performed the experiments, analyzed the data, and prepared the draft manuscript. D. C. Woo performed an MR imaging experiment, analyzed the MR imaging data, and discussed them. Y. J. Chae supported analyzing the MR data. I. Maeng performed THz spectroscopic imaging and supported analyzing THz data. S. Cha analyzed the histological data and added its clinical significance. S. J. Oh and J. Y. Jeon analyzed the data, coordinated the research, acquired research funding, and supervised the project. All authors have reviewed the results and approved the final version of the manuscript. The illustrations used in the figures were created by H. Cheon with the help of the Medical Contents Center of Asan Medical Center.

